# Reprogramming an RNA-guided archaeal TnpB endonuclease for genome editing

**DOI:** 10.1101/2023.03.06.531077

**Authors:** Ying Xu, Tao Liu, Jing Wang, Binyang Xiong, Ling Liu, Nan Peng

## Abstract

RNA-guided endonucleases developed for genome editing have been identified exclusively from bacteria until now. Recently, the RNA-guided TnpB endonucleases encoded by bacterial IS200/IS605 and IS607 families have been reprogrammed for genome editing. In fact, these insertion sequence families are also widely distributed in archaea. However, whether archaea-derived TnpB could be used for genome editing remains unclear. Here, we found that TnpB derived from *Sulfolobus islandicus* REY15A (SisTnpB1) was an Mg^2+^/Mn^2+^-dependent RNA-guided endonuclease, and it could cleave DNA next to the 5’ TTTAA transposon-associated motif (TAM) to generate double-stranded DNA breaks at the 3’-end. We also found that SisTnpB1 was active at 37-85°C and tolerant to variations of TAM. Moreover, SisTnpB1 could be reprogrammed to edit bacterial cells with an editing efficiency of 100% at 37°C or 45°C. In summary, our finding expanded the resources of RNA-guided endonucleases for genome editing, especially those derived from archaea, the third domain of life.

## Introduction

CRISPR-Cas (Clustered regularly interspaced short palindromic repeats and CRISPR-associated) system is prokaryotic immune system, and it can protect bacteria and archaea against invasive viruses and plasmids ^1,2^. The CRISPR ribonuclease (RNP) recognizes DNA or RNA target through a mature CRISPR RNA and cleaves the target in a sequence-specific manner ^3^. The discovery of the CRISPR–Cas system has promoted the development of a new genome editing approach by nucleotide base-pairing. For example, Type II-A and V-A systems use the endonuclease Cas9 ^4^ or Cas12a (also known as Cpf1) ^5^ to specifically pair and cleave the complementary and non-complementary DNA strands through single guide RNA. Cas9 derived from *Streptococcus pyogenes* ^6–9^ and Cas12a derived from *Lachnospiraceae bacterium* and *Acidaminococcus* sp ^5^ have been applied for genome editing. Subsequently, more Cas9 variants have been applied for genome editing, including those from *Staphylococcus aureus* ^10^, *Streptococcus thermophilus* ^11^, *Neisseria meningitidis* ^12,13^, *Campylobacter jejuni* ^14^, and other bacterial species.

CRISPR-Cas9-based innovations have enabled efficient genome engineering in live cells and organisms. However, *in vivo* applications of some Cas9 endonucleases are hindered by their large size, off-target editing, and complex protospacer-adjacent motifs (PAMs). Although small Cas9 proteins such as *S. aureus* Cas9 (1,053 amino acids, 3.16 kbp) have been identified and applied for genome engineering ^10,13,14^, they still exhibit some limitations due to small selection range of PAM sequences and low editing and packing efficiencies. Recently, new programmable endonucleases have been characterized to enrich the genome engineering tools. The potential ancestors of Cas9 and Cas12 family proteins, IscB and TnpB, respectively, have been identified from the widespread IS200/IS605 and IS607 transposon families. Both IscB and TnpB use a single noncoding RNA for RNA-guided cleavage of double-stranded DNA ^15,16^, exhibiting genome editing activity in human cells ^16^.

CRISPR-Cas systems are much more prevalent in archaea, especially in hyperthermophilic archaea, than they are in bacteria ^3^. However, almost all the archaea lack class 2 CRISPR–Cas systems ^3^, except for the identification of one archaeal Cas9 from uncultured microbes ^17^, which limits the discovery of new genome editing nucleases from archaea domain. Considering that TnpB is likely to be the ancestor of Cas12a ^16^ and CasX ^18^, and that transposon families IS200/IS605 encoding TnpB protein are widely present in thermophilic archaea including Sulfolobales ^19^, we investigated the functions of TnpB from *Sulfolobus islandicus* (SisTnpB1) in this study. Our results demonstrated that SisTnpB1 was an RNA-guided thermo-stable endonuclease and was successfully reprogrammed for genome editing in bacterial cells.

## Materials and Methods

### Strains, culture conditions, and transformation

*S. islandicus* REY15A was cultured in SCV medium at 78°C ^20^. *Pediococcus acidilactici* LA412 was cultured in the modified MRS medium at 37°C ^21^. *P. acidilactici* competent cells were prepared and electroporated, as described previously ^21^. *E. coli* DH5α was used for DNA cloning. All *E. coli* strains were cultured in Luria-Bertani (LB) medium at 37°C, and erythromycin (400 μg/ml) was added when required.

### TnpB identification from *S. islandicus*

The genome of *S. islandicus* REY15A was downloaded from NCBI GenBank (Accession No. NC_017276.1). The IS200/IS605 and IS607 transposon families were identified from *S. islandicus* REY15A genome using ISfinder ^22^. Then, 24 TnpB family proteins were identified from the IS200/IS605 and IS607 transposon families, and their amino acids sequences were used for phylogenetic analysis. These TnpB protein sequences from *S. islandicus* REY15A were aligned to ISDra2_TnpB (AAF10241.1) using ClustalW with the default settings, and based on alignment results, a phylogenic tree was constructed using MEGA7 (version 7.0.26) software by neighbor-joining method ^23^, and iTOL was used for downstream visualization of the phylogenic tree ^24^.

### TAM and RE-RNA (ωRNA) analysis

Transposon-associated motif (TAM) sequence has been reported to match the target site sequence of TnpA-mediated transposon ^16^. The DNA sequences from −100 to +50, relative to start codon of *tnpA* genes associated with *tnpB* genes and solo *tnpB* genes were extracted for alignment. Next, the first five conserved bases were predicted to be the TAM sequences. To predict the RE-RNA (ωRNA), nucleotide sequences within 250 bp ahead to100 bp behind the stop codon of each *tnpB* gene were extracted for alignment. Based on the selected conserved regions, ωRNA scaffold was predicted using the “UNAFold Web Server” with its default settings (http://www.unafold.org/DNA_form.php)^25^. The 20 nt sequences located behind the conserved region at the 3’end of the ωRNA were predicted to be the guide sequences.

### Plasmid construction

The *SistnpB1* gene (SiRe_0632) and the predicated ωRNA-coding DNA were amplified from *S. islandicus* REY15A genomic DNA. The *SistnpB1* gene PCR product was digested with Nde I/Sal I, purified, and cloned into pET30a to obtain plasmid pET30a-*SistnpB1*. Then, the ωRNA-coding DNA was digested with Not I/Xho I and cloned into plasmid pET30a-*SistnpB1*. DNA oligo pairs (synthesized by Sangon Biotech, Shanghai) were annealed to generate target DNA, and obtained target DNA was cloned into pUC19 plasmid. The inverse PCR was conducted to obtain target plasmids and mutation *SistnpB1* gene plasmids using the primers carrying the mutation sequences. The PCR products were purified, treated with Dpn I, treated with recombinase, and transformed into *E. coli* DH5α cells. The fluorescence oligos (synthesized by Sangon Biotech, Shanghai) were annealed in a thermo-cycler to generate linear DNA targets for subsequent *in vitro* cleavage. *SistnpB1* gene and ωRNA-coding DNAs respectively under control of the *lacZ* gene promoter and P32 promoter were cloned into pMG36e plasmid through T5 exonuclease DNA assembly (TEDA) to obtain plasmid pSisTnpB1-ωRNA. Then, the DNA donors upstream and downstream target sites were PCR amplified from pMG36e-C-LR ^21^ and cloned into plasmid pSisTnpB1-ωRNA to generate the genome editing plasmid for gene deletion. All primers used in this study were presented in Supplementary Table 1.

### Expression and purification of SisTnpB1 RNP complex

*E. coli* BL21-AI cells carrying *SistnpB1*-ωRNA expression cassette were culture in LB medium supplemented with ampicillin (100 μg/ml) and chloramphenicol (50 μg/ml) at 37°C. After culture to an OD_600_ of 0.6–0.8, protein expression was induced with 0.2% arabinose, and the cells were cultured overnight at 16°C. Then, the cells were harvested by centrifugation, resuspended in 20 mM Tris-HCl (pH 8.0) buffer (containing 250 mM NaCl, 5 mM 2-mercaptoethanol, 25 mM imidazole, 2 mM PMSF, and 5% (v/v) glycerol), and disrupted through high pressure. After removing cell debris by centrifugation, the supernatant was loaded onto the Ni^2+^-charged HiTrap chelating HP column (Cytiva, Marlborough, MA, USA). Subsequently, proteins were eluted with a gradient imidazole (concentration increasing from 25 mM to 500 mM) in the buffer containing 20 mM Tris-HCl (pH 8.0), 500 mM NaCl, 5 mM 2-mercaptoethanol, and 5% (v/v) glycerol. The protein fractions containing SisTnpB1 RNP or the mutant proteins were collected and separated through a Superdex 200 Increase 10/300 GL column (Cytiva, Marlborough, MA, USA), and the peaks of target proteins were analysed by SDS-PAGE. The samples containing SisTnpB1 RNP were dialysed in 20 mM Tris-HCl buffer (pH 8.0 at 25°C) and stored at −80°C or immediately used.

### DNA cleavage assay

Plasmid DNA and synthetic oligoduplex cleavages experiments were conducted at different temperatures (37-85°C) for 60 minutes by adding 5 nM SisTnpB1 RNP complex and 3 μM plasmid DNA carrying different target sequences and TAM sequences in 10 mM Tris-HCl buffer (pH7.5) containing 1 mM DTT, 1 mM EDTA,100 mM NaCl, with 10 mM MgCl_2_, MnCl_2_, CaCl_2_, ZnCl_2_ or NiCl_2_ added into buffer for different reactions. The reaction was stopped by adding 20 mM protease K and 4% SDS solution, and then reaction system was incubated at 37°C for 1 hour. Then, 2 × loading dye was added, and cleavage products were subjected to agarose gel analysis or denaturing PAGE electrophoresis. DNA fragments in agarose gel were visualized by ethidium bromide staining, and DNA fragments in denaturing PAGE were detected using a FUJIFILM scanner (FLA-5100).

### Genome editing in *P. acidilactici* cells

*P. acidilactici* colonies carrying the editing plasmids on the plates containing 5 μg/mL erythromycin were selected randomly, transferred to liquid medium containing 5 μg/mL erythromycin, and incubated at 37°C or 45°C for 12 h. These cell cultures were transferred into fresh medium containing 5 μg/mL erythromycin for the second-round incubation. The editing region was PCR amplified, and PCR products were separated on 1.5% agarose gel. The samples containing shorter bands (shown on gel, representing the successful region deletion) were subjected to DNA sequencing.

## Results

### Identification of TnpB endonucleases from *S. islandicus*

The IS200/IS605 transposon family-encoded TnpB, the potential ancestor of Cas12 and CasX, is widely present in Sulfolobales ^19^. In this study, we found that both typical and short IS200/IS605 transposons encoded *tnpA* and *tnpB* genes or encoded *tnpB* gene alone in *S. islandicus* REY15A (Fig. 1a). To explore the evolution relationships among these TnpB proteins, we constructed a phylogenetic tree using neighbor-joining method. The resulting phylogenetic tree showed that TnpB associated with TnpA, solo TnpB, and TnpB from *Deinococcus radiodurans* ISDra2 ^16^ formed 3 distinct branches, suggesting their evolutionary difference among the 3 types of TnpB proteins (Fig. 1b). Moreover, the tree also showed the relatively close evolutionary relationship between TnpB proteins associated with TnpA and solo TnpB (Fig. 1b). Considering that the transposon/target-associated motif (TAM) recognized by ISDra2 TnpB is identical to the left end (LE) cleavage site-associated motif in *Deinococcus radiodurans* ISDra2 ^16^, we speculated that the TAM sequence recognized by TnpBs might be identical to the LE cleavage site-associated motif in *S. islandicus*. Based on this speculation, we extracted and aligned all the LE sequences of the IS200/IS605 transposons (Fig. S1). We identified conserved DNA sequences at the 5’end of LE (Fig. S1) including a conserved AT-rich TAM sequence (5’-TTTAA-3’) of the IS200/IS605 transposons carrying both *tpnA* and *tnpB* genes (Fig. 1C; Fig. S1a). We also extracted and aligned all the RE sequences encoding ωRNA in *D. radiodurans* ISDra2 IS200/IS605 transposon ^16^. We found that the RE sequences of the transposons encoding both *tpnA* and *tpnB* genes were conserved except their 3’-end sequences (Fig. S2; Fig. 1d). After the transcription, these conserved RE sequences containing two inverted repeats close to the 3’-end (Fig. 1d) probably formed two hairpin motifs as the part of ωRNA scaffold (Fig. 1e). The non-conserved sequences behind the scaffold were predicted to be the guide sequences (Fig. 1d and e). We cloned SiRe_0632 *tnpB* gene and its ωRNA coding sequence into *E. coli* pET30a vector and replaced the putative guide sequence of ωRNA coding sequence with a sequence matching the target DNA. The expression of SiRe_0632 TnpB (SisTnpB1) was induced by IPTG, and then this protein complex was purified from the supernatant of the cell lysate by Ni-NTA column. Then, the Ni-NTA column-purified sample was separated on a gel filtration column, and the sample containing both protein and nucleic acid was subjected to SDS-PAGE analysis (Fig. 1f). SDS-PAGE analysis showed a single band consistent with the molecular weight of SisTnpB1, indicating that nucleic acid was bound to SisTnpB1(Fig. 1f).

**Figure 1.**
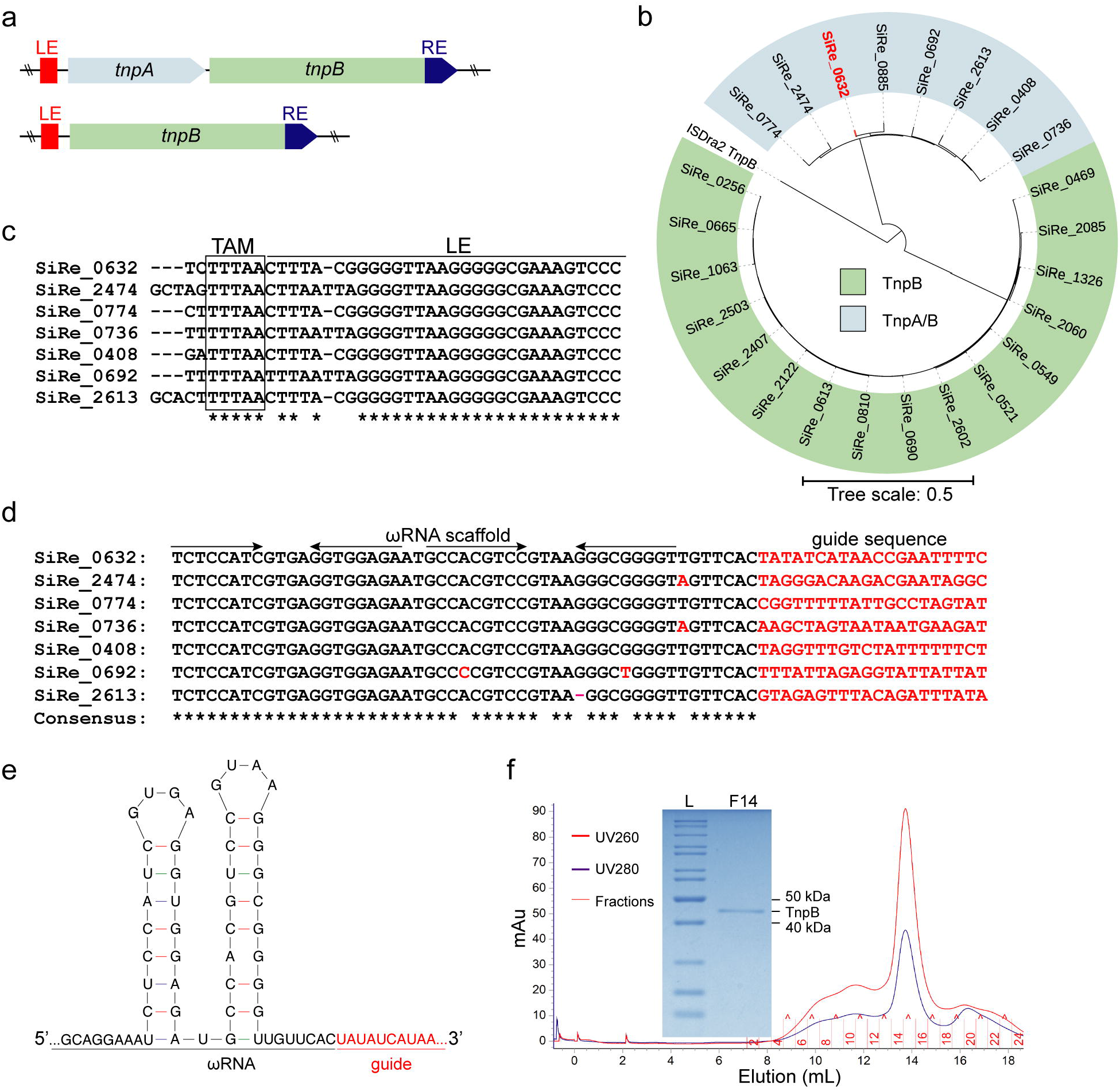
Identification of TnpB endonucleases from *S. islandicus* REY15A. **(a)** Two types of *tnpB* genes, *tnpB* associated with *tnpA* and solo *tnpB*, were identified from *S. islandicus* REY15A. LE, left element flanking the transposon; RE, right element flanking the transposon. **(b)** Phylogenetic tree of TnpB associated with TnpA, solo TnpB, and the TnpB from *D. radiodurans* ISDra2. **(c)** Predicated TAM sequence of *tnpB* genes associated with *tnpA* through alignment. TAM sequences are boxed. **(d)** Alignment of putative ωRNA of *tnpB* genes associated with *tnpA*. (**e**) Predicted structure of ωRNA from the RE of SiRe_0632 *tnpB* gene. (**f**) Purification of SisTnpB1 RNP from *E. coli*. Gel filtration profile of SisTnpB1 RNP and SDS-PAGE analysis of fraction 14 from gel filtration. L: protein ladder; F14, fraction 14.

### SisTnpB1 is a TAM-dependent RNA-guided thermophilic endonuclease

Firstly, we tested the endonuclease activity of SisTnpB1 at different temperatures using plasmid DNA carrying the target DNA sequence with a TAM at its 5’-end (5’-TTTAA-3’). We found that SisTnpB1 cleaved the supercoiled (SC) plasmid DNA into full-length linear (FLL) DNA and open-circle (OC) plasmid within a broad range of temperatures, favoring 65–75°C (Fig. 2a), which was also the optimal growth temperature for *S. islandicus*. Quantitative analysis of the optical density of the product band showed that 23% and 62% FLL DNA products were produced at 37°C and 75°C, respectively (Fig. 2b). Additionally, we found that SisTnpB1 converted supercoiled plasmid DNA into FLL DNA and OC plasmid only in presence of the TAM and target sequence (Fig. 2c). Moreover, SisTnpB1 cleaved plasmid DNA in the presence of Mg^2+^, Mn^2+^, or Ca^2+^ at 37°C with Mn^2+^ being the most effective metal ion (Fig. 2d).

**Figure 2.**
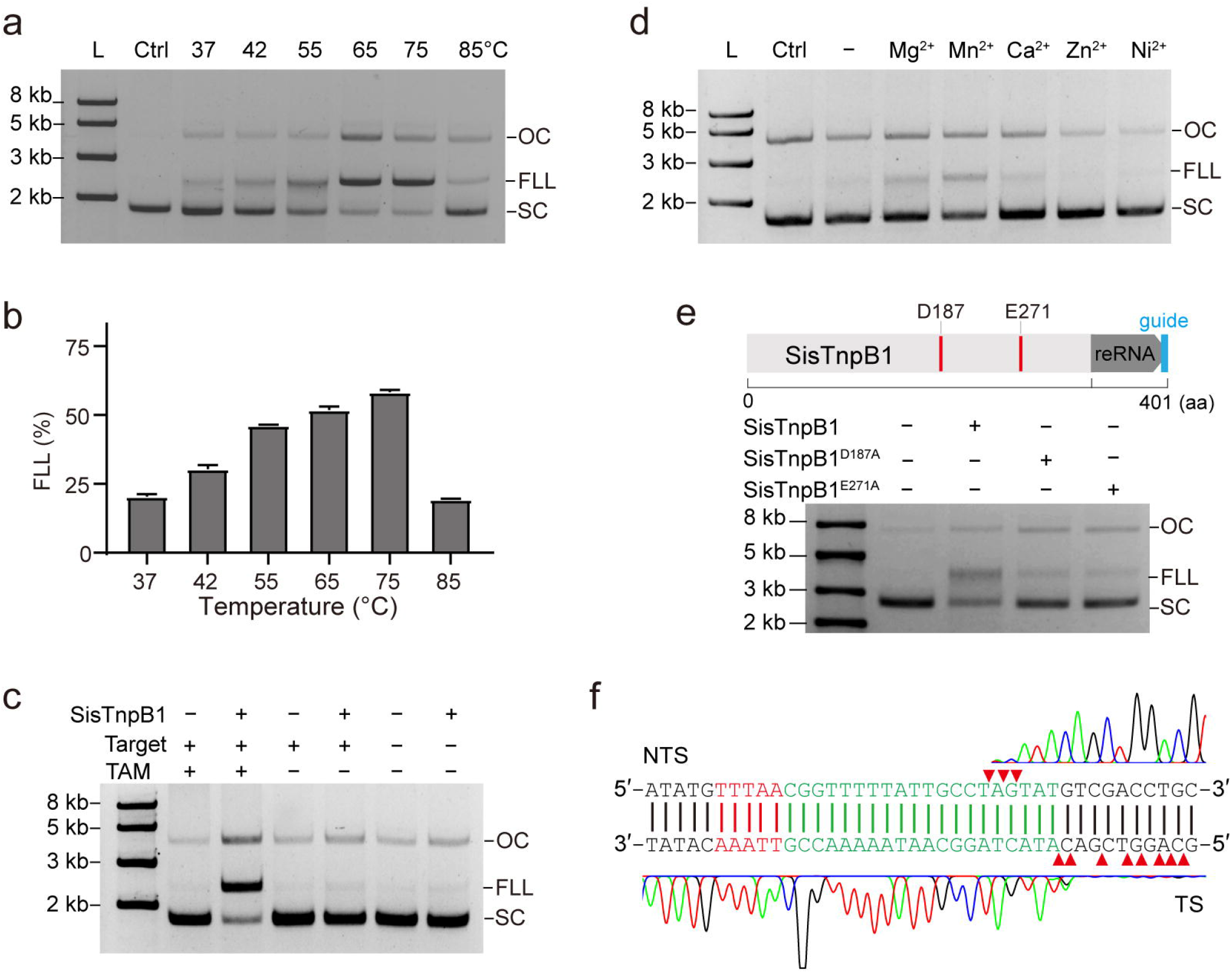
SisTnpB1 is an RNA-guided DNA endonuclease. **(a)** Agarose gel analysis of plasmid DNAs cleaved by SisTnpB1 RNP at different reaction temperature. Plasmids carrying a 5’-TTTAA-3’ TAM and guide RNA-matching 20-bp target DNA sequence was used as cleavage substrate. OC, open-circle; FLL, full-length linear; SC, supercoiled. **(b)** Quantitative analysis of full-length linear cleavage product by SisTnpB1 at different temperatures. **(c)** Plasmid cleavage by the SisTnpB1 RNP complex in presence or absence of TAM or target DNA sequence. **(d)** metal ion-dependent endonuclease activity of SisTnpB1. **(e)** Plasmid cleavage by RNP complexes containing SisTnpB1 mutants generated from mutation of two putative catalytic residues (D187 and E271) in two predictive RuvC domains. **(f)** Run-off sequencing of plasmid products cleaved by SisTnpB1. The red triangles indicate cleavage sites on both targeted strand (TS) and non-targeted strand (NTS).

Two predicated RuvC-like active sites (D187 and E271) were found in the SisTnpB1 amino acid sequence (Fig. 2e). Mutation at either D187 or E271 RuvC-like active site strongly compromised cleavage (Fig. 2e), indicating that the RuvC-like motifs were responsible for cleavage of dsDNA. Run-off sequencing of cleavage products showed a staggered cleavage pattern at 15-18 nt from the TAM on the non-targeting sequence and at 20-28 nt from the TAM on the targeting sequence (Fig. 2f), resulting in 5’ overhangs. Taken together, these results indicate that SisTnpB1 is a TAM-dependent RNA-guided thermophilic endonuclease.

### SisTnpB1 cleaves both dsDNA and ssDNA

We used short double-stranded or single-stranded oligonucleotides with or without TAM and target sequence as the substrates to explore whether SisTnpB1 could cleave both dsDNA and ssDNA. The results showed that SisTnpB1 cleaved dsDNA carrying with both TAM and 20-bp target sequence at 75°C (Fig. 3a and b). PAGE analysis showed the target strand (TS) was completely degraded into fragments (Fig. 3a), while the non-target strand (NTS) was only partially cleaved (Fig. 3b). At 37°C, the cleavage of both TS and NTS sequences was dramatically reduced (Fig. 3c and d). SisTnpB1 showed no cleavage activity at the absence of the ωRNA matching sequence at 75°C (Fig. S3a and b). SisTnpB1 showed very weak cleavage activity at the target sequence strand, no cleavage at the non-target strand of the dsDNA substrate at the absence of a TAM sequence (Fig. 3e and f) at 75°C, and almost no cleavage activity at 37°C (Fig. 3g and h). SisTnpB1 also cleaved a matched single-stranded DNA at 75°C or 37 °C with or without the presence of TAM (Fig. 3i - l). In the presence of a TAM sequence in the ssDNA substrate resulted in high cleavage efficiency (Fig. 3i and k). Lastly, SisTnpB1 showed no cleavage activity on the dsDNA (Fig. S3a and b) or ssDNA without target DNA sequence (Fig. S3c and d) at 75°C.

**Figure 3.**
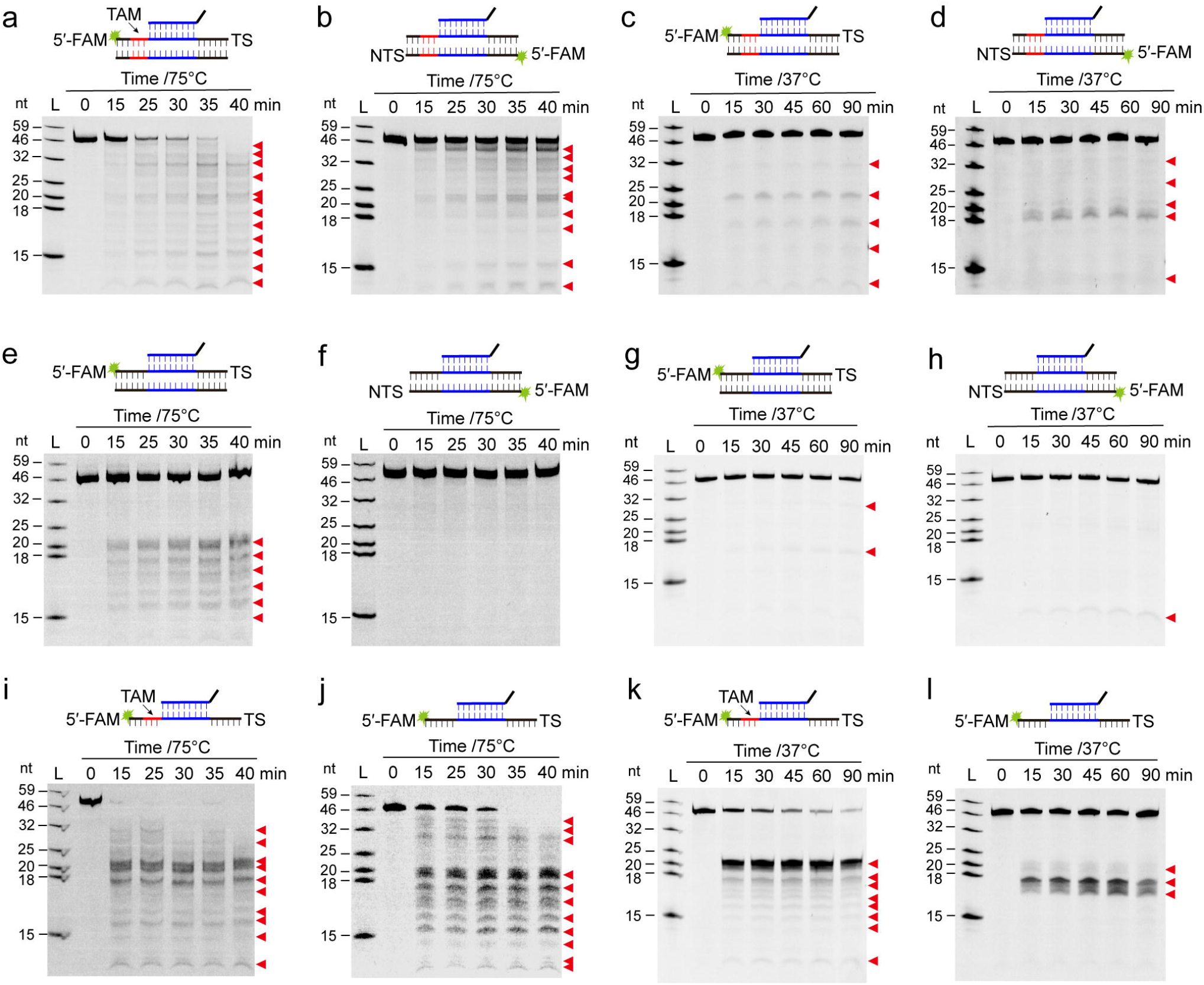
SisTnpB1 cleaves both dsDNA and ssDNA. (**a - d**) 20% denaturing PAGE analysis of 59-bp target dsDNA carrying TAM cleaved by SisTnpB1 RNP at 75°C **(a** and **b)** or 37°C **(c** and **d)**. Target strands **(a** and **c)** or non-target strands **(b** and **d)** were FAM-labelled at 5’ends. **(e - h)** 20% denaturing PAGE analysis of 59-bp target dsDNA carrying no TAM cleaved by SisTnpB1 RNP at 75°C **(e** and **f)** or 37°C **(g** and **h)**. **(I - l)** 20% denaturing PAGE analysis of SisTnpB1 RNP-cleaved 59 nt non-target ssDNA carrying TAM **(I** and **k)** or without TAM **(j** and **l)** at 75°C **(i** and **j)** or 37°C **(k** and **l)**. ssDNAs were FAM-labelled at all 5’ends. Cleaved products were indicated by red triangles.

### Effects of seed and TAM sequence variation on SisTnpB1 endonuclease activity

Transversion mutations (A ↔ T or G ↔ C) were introduced into the target sequence and the adjacent TAM sequence (Fig. 4a) to investigate the effect of seed and TAM sequences on SisTnpB1 endonuclease activity. Mutation M1~5 represented transversion of + 1GCCAA+5 into +1CGGTT+5 on the target sequence, and this mutation almost abolished DNA cleavage by SisTnpB1 (Fig. 4b). Similarly, mutation M6~10 drastically reduced DNA cleavage, and M11~15 and M16~20 had weak effect on target DNA cleavage (Fig. 4b), indicating that the seed sequence was located at the site from +1 to +10. Moreover, M-1~-5 with transversion mutation of the TAM sequence almost eliminated DNA cleavage by SisTnpB1 (Fig. 4b), suggesting the importance of TAM sequence for target DNA cleavage by SisTnpB1. Then, we introduced single transversion mutations to the seed and TAM sequences and found that transversion mutation at +1 nucleotide (C ↔ G) of the target sequence strongly inhibited SisTnpB1 cleavage, while single transversions at other sites had less effect on target DNA cleavage by SisTnpB1 (Fig. 4c). These results indicated that SisTnpB1 was highly tolerant to mutations on the target DNA sequence. Saturation mutations were introduced to TAM to examine the effects of TAM variations on SisTnpB1 cleavage. The results showed that transversion and transition mutations (such as A to T, G to C) at any nucleotide on the TAM sequence had weak effect on the target DNA cleavage by SisTnpB1 (Fig. 4d), except for the mutant M-5 (T to C), M-5 (T to G) and M-3(T to G), which showed less than 20% cleavage efficiency at 75°C (Fig. 4e). This result indicated that SisTnpB1 required a less conserved TAM sequence (5’-WNHNN-3’) for recognizing target DNA, and thus it was capable of targeting DNA with a broad range of TAM sequences.

**Figure 4.**
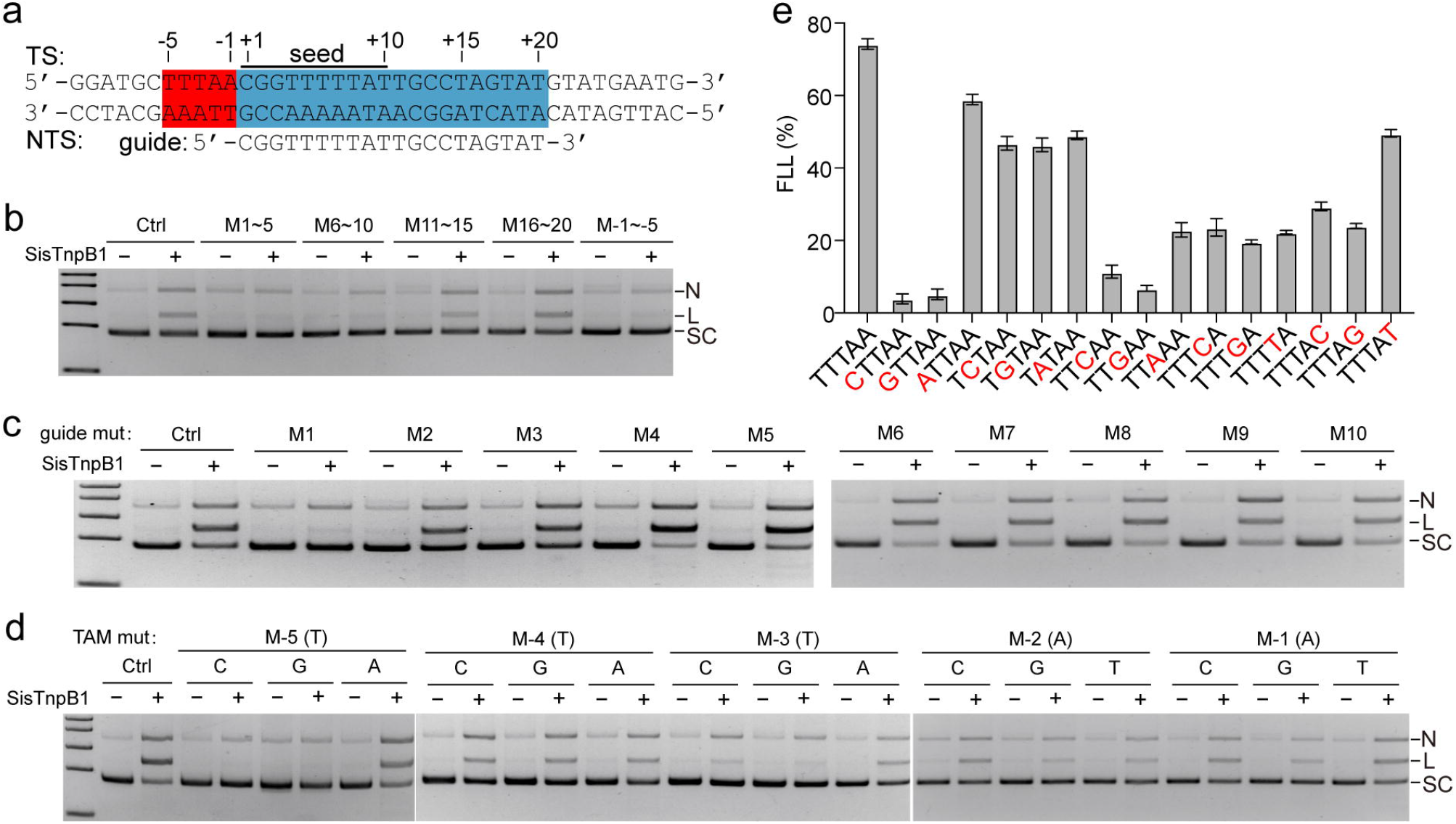
Effects of seed and TAM sequence variation on SisTnpB1 endonuclease activity. **(a)** Schematic of the target dsDNA sequence in the plasmid and guide RNA sequence of the ωRNA. TAM and guide-matching sequences were boxed by red and light blue rectangles, respectively. The location of bases in the TAM and target sequences were labeled by numbers. **(b)** Agarose gel analysis of SisTnpB1 RNP-cleaved products from the substrate plasmid with target DNA sequence or TAM sequence mutations. Numbers indicated the mutation regions. **(c** and **d)** Agarose gel analysis of the SisTnpB1 RNP-cleaved products from the substrate plasmid with screen mutations at target DNA sequence **(c)** or with saturation mutations at the TAM sequence **(d)**. **(e)** Quantitative analysis of full-length linear SisTnpB1-cleaved product from the substrate plasmid with mutated TAM sequences.

### SisTnpB is reprogrammed for *in vivo* DNA targeting and genome editing in bacterial cells

*P. acidilactici* LA412 grew at condition of the temperature from 37°C to 55°C well (Fig. S4). Therefore, it was a very suitable host to investigate the functions of the thermophilic programmable DNA endonuclease *in vivo*. Firstly, we examined target DNA cleavage efficiency by SisTnpB1 in *P. acidilactici* LA412 carrying two endogenous plasmids ^21^. We selected a 20-bp target sequence with a 5’-end TTTAA sequence, and this 20-bp target sequence was present in both GE00014 gene of plasmid 1 and GE00039 gene of plasmid 2. We used this target sequence as the guide of the ωRNA and cloned ωRNA sequence and *SistnpB1* gene into the plasmid pMG36e to obtain an interference plasmid (Fig. 5a). The transformants carrying the interference plasmid were cultured on the plates at 37°C. The target region on GE00014 gene on plasmid 1 and GE00039 gene on plasmid 2, or GE00037 gene on plasmid 1 and GE00033 gene on plasmid 2 of these transformants was PCR amplified. The agarose gel analysis showed that the bands of PCR products on were weaker than those of the wildtype control (Fig. 5b). However, after one-round passage of cells in modified MRS medium containing 5 μg/mL erythromycin, no bands of PCR products of the target regions were detected (Fig. 5B), indicating that the endogenous plasmids were depleted through *in vivo* DNA cleavage by the plasmid-encoded SisTnpB1 RNP.

**Figure 5.**
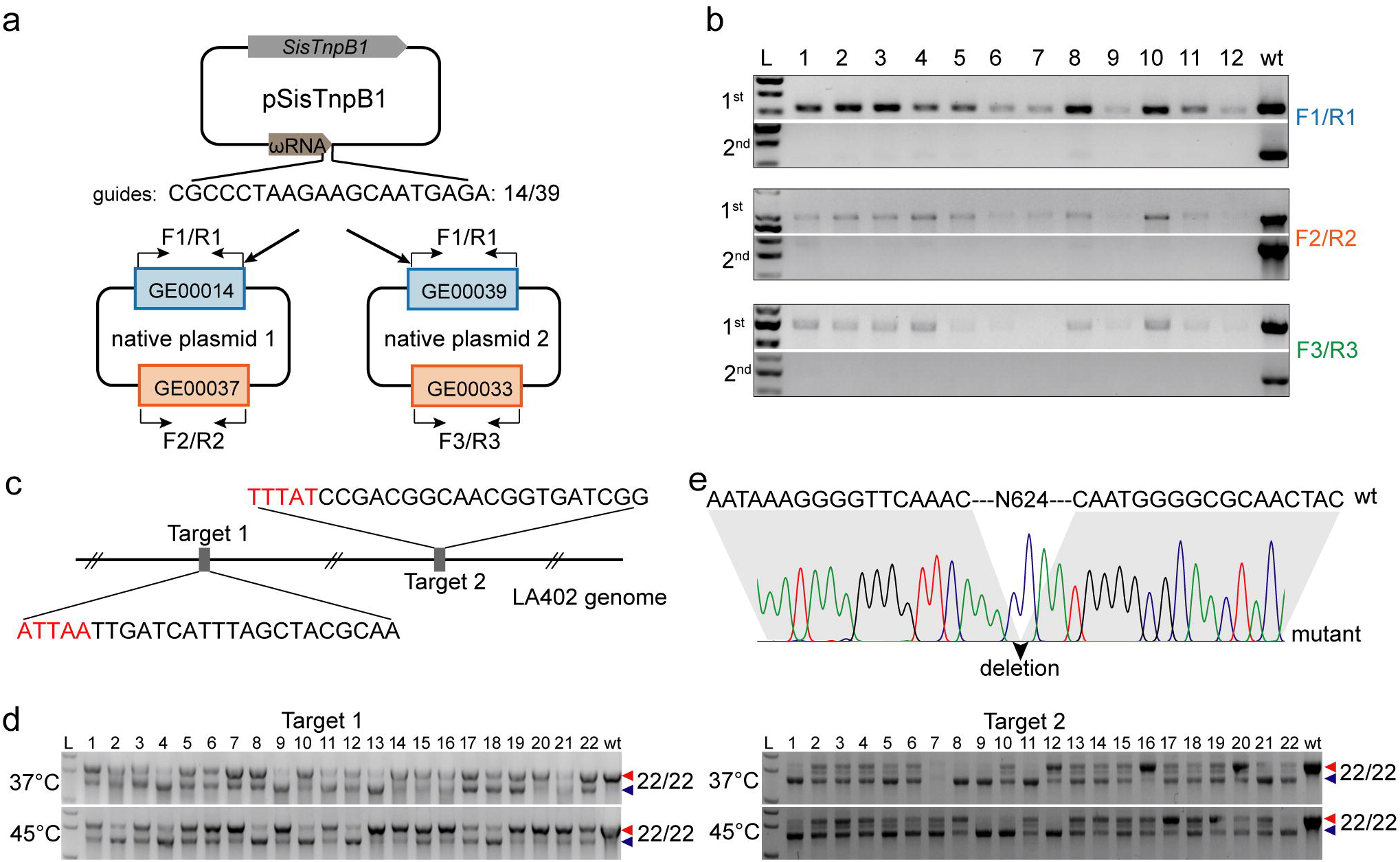
Plasmid interference and genome editing in bacterial cells. **(a)** Schematic of pMG36e-based plasmid interference. Interference plasmid encodes *SistnpB1* gene and the ωRNA, thus targeting GE00014 of endogenous plasmid 1 and GE00039 gene of endogenous plasmid 2. Primer pairs F1/R1, F2/R2, and F3/R3 were used to amplify GE00014/GE00039 gene, GE00037 gene, and GE00033 gene, respectively. **(b)** Agarose gel analysis of PCR products from randomly selected colonies (1^st^) or those after one-round passage (2^nd^). The primer pairs F1/R1, F2/R2, and F3/R3 were used for PCR amplification. **(c)** Selected targets in *pyrE* gene of *P. acidilactici* LA412 genome. Black letters indicate the target sequences, and red letters indicate TAM sequences. **(d)** Agarose gel analysis of PCR products from randomly selected transformants after one round of passage at 37°C or 45°C. Red and blue triangles indicate wildtype or deletion bands. L, DNA ladder; 1-22, 22 randomly selected transformants; wt: wildtype cell control. (**e**) DNA sequencing of PCR products from the *pyrE* deletion mutant carrying the editing plasmid targeting site 1. The black triangle indicates the site of a 624-bp deletion.

In order to establish the genome editing tool based on SisTnpB1, we selected two target sites respectively adjacent to 5’ ATTAA and 5’ TTTAT motif on *pyrE* gene (Fig. 5c). The pMG36e-based editing plasmids encoding SisTnpB1 and the ωRNA with a guide matching to target site 1 or 2 on *pyrE* gene were electroporated into *P. acidilactici* LA412 cells to obtain transformants, and transformants on the plates were randomly selected and incubated in the liquid medium containing 5 μg/mL erythromycin. The target regions were PCR amplified, and the agarose gel analysis indicated that at 37°C and 45°C, genome editing efficiency at target site 1 of the transformants carrying desired gene deletion was 36.4% and 54.5%, and that at target site 2 was 95.5% and 90.9%, respectively (Fig. S5). After one-round passage of the transformants in the fresh MRS medium containing 5 μg/mL erythromycin, the genome editing efficiency increased to 100% for both target site 1 and 2 at 37°C and 45°C (Fig. 5d). The alignment of DNA sequencing results with wildtype genome sequence verified a 624 bp deletion on *pyrE* gene, indicating a satisfactory genome editing (Fig. 5e). The above results suggested that SisTnpB1 could be used for genome editing in the hosts living at 37°C or higher temperature.

## Discussion

Insertion sequences from the IS200/IS605 and IS607 families are among the simplest and most ancient mobile genetic elements (MGEs) ^26^, and they typically encode two open reading frames (ORFs) transposase gene *tnpA* and *tnpB*, and carry subterminal LE and RE palindromic elements ^26^. The *tnpB* gene is not required for transposition ^27^ but it can reduce the transposition activity ^28^. Recently, TnpB protein has been identified as RNA-guided endonucleases in *Deinococcus radiodurans* ISDra2 ^16^, *Ktedonobacter racemifer* strain SOSP1-21 and *Alicyclobacillus macrosporangiidus* ^15^. Moreover, ISDra2 TnpB has been reprogrammed to edit human cells ^16^. Although the IS200/IS605 and IS607 families in bacteria have been well studied to date ^29^, little is known about their presence in archaeal genomes ^19^. IS200/IS605 and IS607 families are present in hyperthermophiles Sulfolobales/Thermoproteales/Thermococcales/Thermoplasmales, halophiles Halobacteriales, and methanogens Methamosarcinales ^19^. Therefore, studying IS200/IS605 and IS607 families, especially those programmable endonucleases from archaea domain will provide more probable genome editing tools for multi-scenario application.

The evolution of CRISPR-Cas9 from IscB and CRISPR-Cas12 from TnpB ^15^ will lead to identification of more ancestral programmable endonucleases from MGEs. The Cas nucleases and their ancestors showed similar reaction pattern that both require a guide RNA and a PAM or TAM sequence for target recognition. Cas12 nucleases use a guide RNA originated from the CRISPR array, whereas ISDra2 TnpB, AmaTnpB, and SisTnpB1 use right transposon element-derived reRNA (also known as ωRNA) as a guide RNA^15,16^ (Fig. 1d). Moreover, each Cas12a endonuclease requires a T-rich PAM sequence ^5^. However, the smaller Cas12f (also known as Cas14) is a ssDNA-targeting CRISPR endonuclease, and it requires no PAM sequence ^30^. Similarly, ISDra2 TnpB and AmaTnpB recognize 5’-end TTGAT TAM sequence and TCAC TAM sequence, respectively ^15,16^. In this study, we identified an RNA-guided TnpB endonuclease from *S. islandicus*, and it had a wide range of temperature adaptability and thermo-stability (Fig. 2a and b). Besides, no non-specific dsDNA cleavage by SisTnpB1 was detected in our study (Fig. S3a and b). Our identification of programmable DNA nucleases from archaea expanded genome editing toolbox. The *in vitro* experiment confirmed that SisTnpB1 recognized a broad range of TAM sequences 5’-WNHNN-3’ (Fig. 4d and e), and the genome editing experiments showed that SisTnpB1 had a high capacity for recognition and cleavage of a given DNA sequence in *P. acidilactici* cells (Fig. 5d and e).

Programmable endonucleases from thermophiles are required as the ribonucleo-proteins (RNPs) since they are very stable and easy to be delivered into the target tissue or bloodstream of the patients or organisms ^31^. They are also required in producing biofuels and biochemicals in the emerging metabolic platforms and engineering extreme thermophiles ^32^. Recently, Thermo-stable Cas9 and Cas12 have been identified from bacteria. For example, thermo-stable Cas9 proteins have been identified from *Geobacillus* ^33,34^ and *Ignavibacterium* ^35^. Cas9 proteins from *Geobacillus* exhibit programable DNA cleavage activity below 70°C with the maximum cleavage activity at ~60°C ^33,34^, while IgnaviCas9 display activity at the temperature up to 100°C ^35^. Meanwhile, thermostable Cas12b proteins (with activity at 70°C) from *Brevibacillus* ^36,37^ have been characterized and used for one-pot detection of SARS-CoV-2 variants ^36^. The optimal reaction temperature for all these CRISPR endonucleases is below 70°C, except for IgnaviCas9 which has a wide optimal temperature ranging from 31°C to 100°C. However, IgnaviCas9 is found to be less active even at the optimal temperature ^35^. Our data showed that SisTnpB1 was active even at 85°C (Fig. 2a and b), and it could completely degrade the target DNA at 75°C (Fig. 3a). Importantly, SisTnpB1 exhibited an editing efficiency up to100% in bacterial cells at 37°C or higher temperature (Fig. 5d), suggesting its potential broad application for different organisms living at different temperatures. Moreover, SisTnpB1 had only 401 amino acid residues, which was much smaller in size than above-mentioned thermostable Cas nucleases. These advantages of SisTnpB1 were critical for new molecular biology applications under harsh conditions such as requirement for the cleavage activity over a wide temperature range.

## Acknowledgements

This work was supported by grants from the National Key Research and Development Program of China (2022YFA0912200), the National Natural Science Foundation of China (No. 32270090), the Foundation of Hubei Hongshan Laboratory (No. 2021hszd013 and 2021hszd022), and the LongYun Program for College of Life Science and Technology, Huazhong Agricultural University. Funding for open access charge: the National Key Research and Development Program of China (2022YFA0912200).

## Competing interests

All authors declare that they have a patent pending to this material.

## Notes

### Summary of Updates

Supplementary files updated; Title revised.

## Rferences

1 Barrangou, R. et al. CRISPR provides acquired resistance against viruses in prokaryotes. Science 315, 1709–1712, doi:10.1126/science.1138140 (2007).

2 Makarova, K. S. et al. Evolution and classification of the CRISPR-Cas systems. Nat Rev Microbiol 9, 467–477, doi:10.1038/nrmicro2577 (2011).

3 Makarova, K. S. et al. Evolutionary classification of CRISPR-Cas systems: a burst of class 2 and derived variants. Nat Rev Microbiol 18, 67–83, doi:10.1038/s41579-019-0299-x (2020).

4 Jinek, M. et al. A programmable dual-RNA-guided DNA endonuclease in adaptive bacterial immunity. Science 337, 816–821, doi:10.1126/science.1225829 (2012).

5 Zetsche, B. et al. Cpf1 is a single RNA-guided endonuclease of a class 2 CRISPR-Cas system. Cell 163, 759–771, doi:10.1016/j.cell.2015.09.038 (2015).

6 Cong, L. et al. Multiplex genome engineering using CRISPR/Cas systems. Science 339, 819–823, doi:10.1126/science.1231143 (2013).

7 Mali, P. et al. RNA-guided human genome engineering via Cas9. Science 339, 823–826, doi:10.1126/science.1232033 (2013).

8 Jinek, M. et al. RNA-programmed genome editing in human cells. eLife 2, e00471, doi:10.7554/eLife.00471 (2013).

9 Jiang, W., Bikard, D., Cox, D., Zhang, F. & Marraffini, L. A. RNA-guided editing of bacterial genomes using CRISPR-Cas systems. Nat Biotechnol 31, 233–239, doi:10.1038/nbt.2508 (2013).

10 Ran, F. A. et al. In vivo genome editing using *Staphylococcus aureus* Cas9. Nature 520, 186–191, doi:10.1038/nature14299 (2015).

11 Esvelt, K. M. et al. Orthogonal Cas9 proteins for RNA-guided gene regulation and editing. Nat Methods 10, 1116–1121, doi:10.1038/nmeth.2681 (2013).

12 Hou, Z.et al. Efficient genome engineering in human pluripotent stem cells using Cas9 from *Neisseria meningitidis*. Proc Natl Acad Sci USA 110, 15644–15649, doi:doi:10.1073/pnas.1313587110 (2013).

13 Edraki, A. et al. A compact, high-accuracy Cas9 with a dinucleotide PAM for *in vivo* genome editing. Mol Cell 73, 714–726 e714, doi:10.1016/j.molcel.2018.12.003 (2019).

14 Kim, E. et al. In vivo genome editing with a small Cas9 orthologue derived from *Campylobacter jejuni*. Nat Commun 8, 14500, doi:10.1038/ncomms14500 (2017).

15 Altae-Tran, H. et al. The widespread IS200/IS605 transposon family encodes diverse programmable RNA-guided endonucleases. Science 374, 57–65, doi:10.1126/science.abj6856 (2021).

16 Karvelis, T. et al. Transposon-associated TnpB is a programmable RNA-guided DNA endonuclease. Nature 599, 692–696, doi:10.1038/s41586-021-04058-1 (2021).

17 Burstein, D. et al. New CRISPR-Cas systems from uncultivated microbes. Nature 542, 237–241, doi:10.1038/nature21059 (2017).

18 Liu, J. J. et al. CasX enzymes comprise a distinct family of RNA-guided genome editors. Nature 566, 218–223, doi:10.1038/s41586-019-0908-x (2019).

19 Filee, J., Siguier, P. & Chandler, M. Insertion sequence diversity in archaea. Microbiol Mol Biol Rev 71, 121–157, doi:10.1128/MMBR.00031-06 (2007).

20 Deng, L., Zhu, H., Chen, Z., Liang, Y. X. & She, Q. Unmarked gene deletion and host-vector system for the hyperthermophilic crenarchaeon *Sulfolobus islandicus*. Extremophiles 13, 735–746, doi:10.1007/s00792-009-0254-2 (2009).

21 Liu, L. et al. High-efficiency genome editing based on endogenous CRISPR-Cas system enhances cell growth and lactic acid production in *Pediococcus acidilactici*. Appl Environ Microbiol 87, e0094821, doi:10.1128/AEM.00948-21 (2021).

22 Siguier, P., Perochon, J., Lestrade, L., Mahillon, J. & Chandler, M. ISfinder: the reference centre for bacterial insertion sequences. Nucleic Acids Res 34, D32–36, doi:10.1093/nar/gkj014 (2006).

23 Kumar, S., Stecher, G. & Tamura, K. MEGA7: Molecular evolutionary genetics analysis version 7.0 for bigger datasets. Mol Biol Evol 33, 1870–1874, doi:10.1093/molbev/msw054 (2016).

24 Letunic, I. & Bork, P. Interactive Tree Of Life (iTOL) v5: an online tool for phylogenetic tree display and annotation. Nucleic Acids Res 49, W293–W296, doi:10.1093/nar/gkab301 (2021).

25 Zuker, M. Mfold web server for nucleic acid folding and hybridization prediction. Nucleic Acids Res 31, 3406–3415, doi:10.1093/nar/gkg595 (2003).

26 Siguier, P., Gourbeyre, E. & Chandler, M. Bacterial insertion sequences: their genomic impact and diversity. FEMS Microbiol Rev 38, 865–891, doi:10.1111/1574-6976.12067 (2014).

27 Kersulyte, D. et al. Transposable element ISHp608 of *Helicobacter pylori:*nonrandom geographic distribution, functional organization, and insertion specificity. J Bacteriol 184, 992–1002, doi:10.1128/jb.184.4.992-1002.2002 (2002).

28 Pasternak, C. et al. ISDra2 transposition in *Deinococcus radiodurans* is downregulated by TnpB. Mol Microbiol 88, 443–455, doi:10.1111/mmi.12194 (2013).

29 Barabas, O. et al. Mechanism of IS200/IS605 family DNA transposases: activation and transposon-directed target site selection. Cell 132, 208–220, doi:10.1016/j.cell.2007.12.029 (2008).

30 Harrington, L. B. et al. Programmed DNA destruction by miniature CRISPR-Cas14 enzymes. Science 362, 839–842, doi:10.1126/science.aav4294 (2018).

31 Staahl, B. T. et al. Efficient genome editing in the mouse brain by local delivery of engineered Cas9 ribonucleoprotein complexes. Nat Biotechnol 35, 431–434, doi:10.1038/nbt.3806 (2017).

32 Crosby, J. R. et al. Extreme thermophiles as emerging metabolic engineering platforms. Curr Opin Biotechnol 59, 55–64, doi:10.1016/j.copbio.2019.02.006 (2019).

33 Harrington, L. B. et al. A thermostable Cas9 with increased lifetime in human plasma. Nat Commun 8, 1424, doi:10.1038/s41467-017-01408-4 (2017).

34 Mougiakos, I. et al. Characterizing a thermostable Cas9 for bacterial genome editing and silencing. Nat Commun 8, 1647, doi:10.1038/s41467-017-01591-4 (2017).

35 Schmidt, S. T., Yu, F. B., Blainey, P. C., May, A. P. & Quake, S. R. Nucleic acid cleavage with a hyperthermophilic Cas9 from an uncultured *Ignavibacterium*. Proc Natl Acad Sci U S A 116, 23100–23105, doi:10.1073/pnas.1904273116 (2019).

36 Nguyen, L. T. et al. A thermostable Cas12b from *Brevibacillus leverages* one-pot discrimination of SARS-CoV-2 variants of concern. EBioMedicine 77, 103926, doi:10.1016/j.ebiom.2022.103926 (2022).

37 Tian, Y. et al. A novel thermal Cas12b from a hot spring bacterium with high target mismatch tolerance and robust DNA cleavage efficiency. Int J Biol Macromol 147, 376–384, doi:10.1016/j.ijbiomac.2020.01.079 (2020).

