## Supplementary Figures and Table for "Reprogramming an RNA-guided archaeal TnpB endonuclease for genome editing"

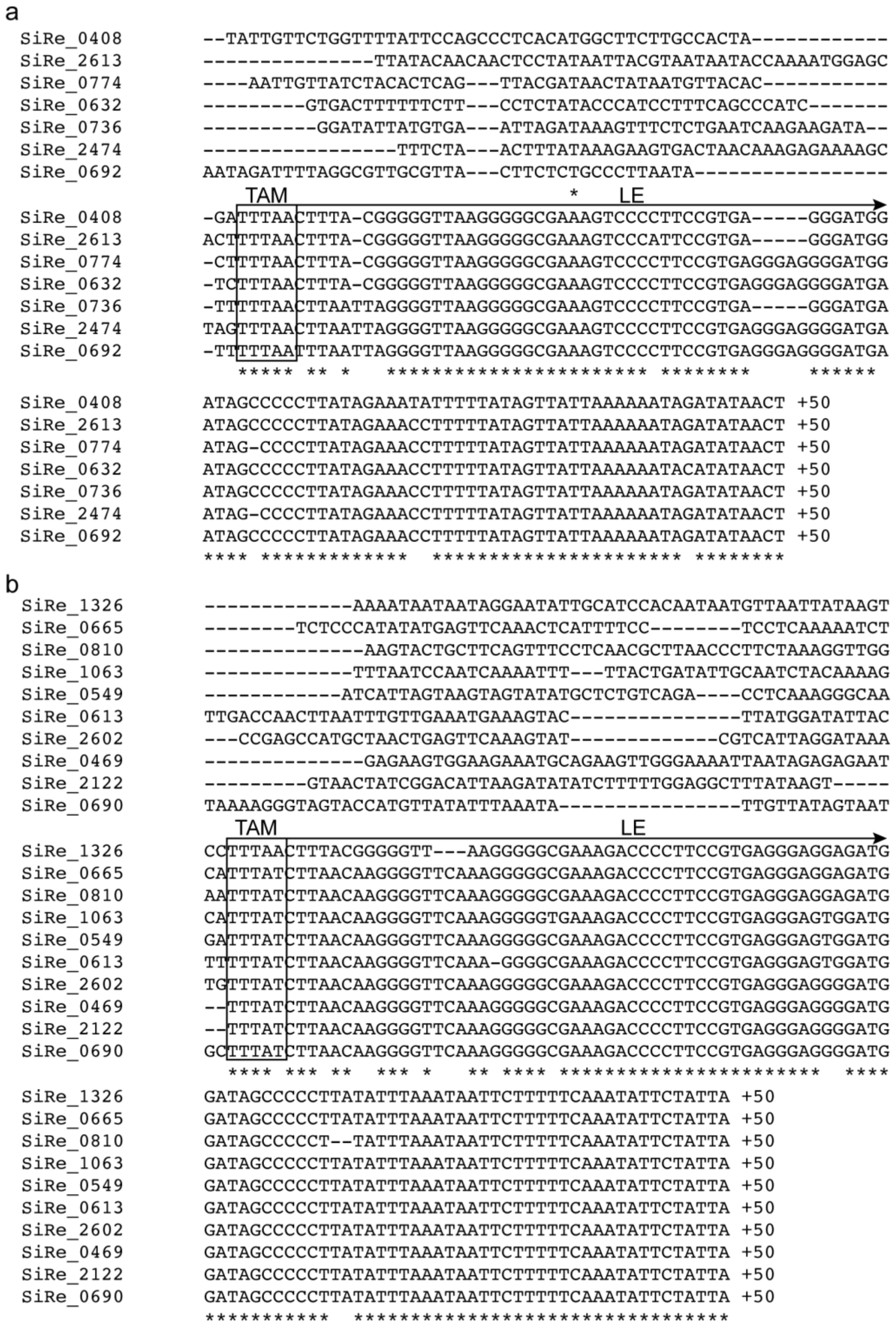


**Supplementary Figure 1. Identification of TAM sequences.** The DNA sequences from -100 to +50 relative to the start codon of eight *tnpA* genes associated with *tnpB* **(a)** and solo *tnpB* genes **(b)** were extracted and analyzed through alignment. The first five conserved bases were predicted to be the TAM sequences. TAM sequences are boxed and LE sequences are indicated.

```
SiRe_0692  TTAAGAAGGATTTGACTTTGGCTGACCGTATATTCTCATGTCCTAAATGTGGTTGGACT
SiRe_0774  TTAAGAAGGACTTGACTTTGGCTGACCGTGTGTTTGTATGTCCTAAATGTGGTTGGACT
SiRe_0632  TTAAGAAGGACTTGGCTTTGGCTGACCGTGTGTTTGTATGCCCTGAGTGCGGTTGGACT
SiRe_2474  TTAAGAAGGACTTGGCTTTGGCTGACCGTGTGTTTGTATGCCCCAAGTGTGGTTGGACT
SiRe_0736  TTAAGAAGGACCTGGCTTTGGCTGACCGTGTGTTTGTATGCCCCAAGTGTGGTTGGGCT
SiRe_2613  TTAAGAAGGATTTGACTTTGGCTGACCGTGTGTTTGTATGTCCCAAGTGTGGTTGGACT
SiRe_0408  TTAAGAAGGACTTGGCTTTGGCTGACCGTGTGTTTGTATGTCCCAAGTGTGGTTGGGCT
           **********  ** ************** * **   *** **  * ** ****** **

SiRe_0692 GTAGATCGTGACTATAATGCTTCTCTAAATATTCTTCATGCGGGGTCGGGACAGCCCTTA
SiRe_0774 GTAGATCGTGACTATAATGCTTCTCTAAATATTCTTCGTGCGGGGTCGGGACTGCCCTTA
SiRe_0632 GTAGATCGTGACTATAATGCTTCTCTAAATATTCTTCGTGCGGGGTCGGGACTGCCCTTA
SiRe_2474 GTAGATCGTGACTATAATGCTTCTCTAAATATTCTTCGTGCGGGGTCGGGACTGCCCTTA
SiRe_0736 GTAGATCGTGACTATAATGCTTCTCTAAATATTCTTCGTGCGGGGTCGGGACAGCCCTTA
SiRe_2613 GTAGACCGTGACTATAATGCTTCTCTAAATATTCTTCATGCGGGGTCGGGACAGCCCTTA
SiRe_0408 GTAGATCGTGACTATAATGCTTCTCTAAATATTCTTCATGCGGGGTCGGGACAGCCCTTA
          ***** ****************************** *************** *******

SiRe_0692 GAGCCTGTGGACAGGAGACCTCTGCTATACATTCCCTTCTCTGAGGGTGTGTATAGTAAG
SiRe_0774 GAGCCTGTGGACAGGGGACCTCTGCTATACATTCCCTTCTCAGAAGGGGTGTATAGTAAG
SiRe_0632 GAGCCTGTGGACAGGGGACCTCTGCTATACATTCCCTTCTCTGAGGGTGTGTATAGTAAG
SiRe_2474 GAGCCTGTGGACAGGGGACCTCTGCTATACATACCCTTCTCTGAGGGTGTGTATAGTAAG
SiRe_0736 GAGCCTGTGGACAGGGGACCTCTGCTATACATACCCTTCTCTGAGGGTGTGTATAGTAGG
SiRe_2613 GAGCCTGTGGACAGGGGACCTCTGCTATACATACCCTTCTCTGAGGGTGTGTATAGTAAG
SiRe_0408 GAGCCTGTGGACAGGGGACCTCTGCTATACATACCCTTCTCTGAGGGTGTGTATAGTAGG
          *************** **************** ******** ** ** ********** *

SiRe_0692 TTTCCTGGAAGAAGCAGGAAATCTCCATCGTGAGGTGGAGATGCCCCGTCCGTAAGGGCT
SiRe_0774 TTTCTTGGAAGAAGCAGGAAATCTCCATCGTGAGGTGGAGATGCCACGTCCGTAAGGGCG
SiRe_0632 TTTCTTGGAAGAAGCAGGAAATCTCCATCGTGAGGTGGAGATGCCACGTCCGTAAGGGCG
SiRe_2474 TTTCTTGGAAGAAGCAGGAAATCTCCATCGTGAGGTGGAGATGCCACGTCCGTAAGGGCG
SiRe_0736 TTTCCTGGAAGAAGCAGGAAATCTCCATCGTGAGGTGGAGATGCCACGTCCGTAAGGGCG
SiRe_2613 TTTCCTGGAAGAAGCAGGAAATCTCCATCGTGAGGTGGAGATGCCACGTCCGTAA-GGCG
SiRe_0408 TTTCCTGGAAGAAGCAGGAAATCTCCATCGTGAGGTGGAGATGCCACGTCCGTAAGGGCG
          **** **************************************** ********* ***

SiRe_0692 GGGTTGTTCACTTTATTAGAGGTATTATTAT
SiRe_0774 GGGTTGTTCACCGGTTTTTATTGCCTAGTAT
SiRe_0632 GGGTTGTTCACTATATCATAACCGAATTTTC
SiRe_2474 GGGTAGTTCACTAGGGACAAGACGAATAGGC
SiRe_0736 GGGTAGTTCACAAGCTAGTAATAATGAAGAT
SiRe_2613 GGGTTGTTCACGTAGAGTTTACAGATTTATA
SiRe_0408 GGGTTGTTCACTAGGTTTGTCTATTTTTTCT
          **** ******     guide sequence
```

**Supplementary Figure 2. Identification of ωRNA sequences.** The ωRNA sequences located at the 3’-end of *tnpB* genes associated with *tnpA* gene were aligned, and the guide sequences were boxed.

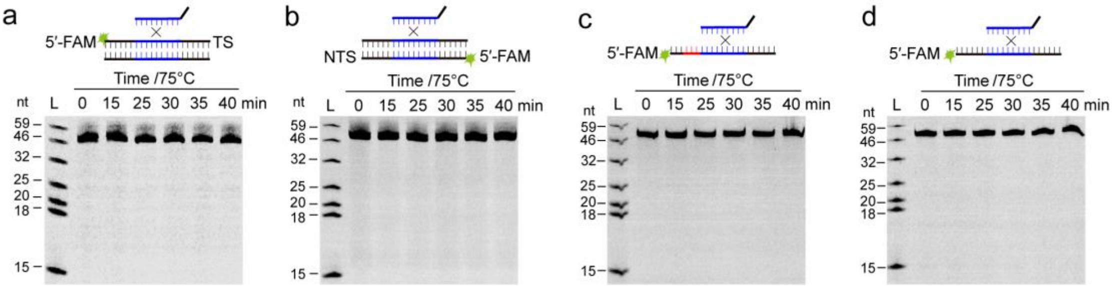


**Supplementary Figure 3. SisTnpB1 RNP shows no non-specific endonuclease activity.** 20% denaturing PAGE analysis of SisTnpB1 RNP-cleaved 59-bp dsDNA (**a** and **b**) or 59-nt ssDNA (**c** and **d**) without matching sequence. Target strands **(a**, **c** and **d)** or non-target strand **(b)** were FAM-labelled at 5’-end. TAM sequence is indicated in red.

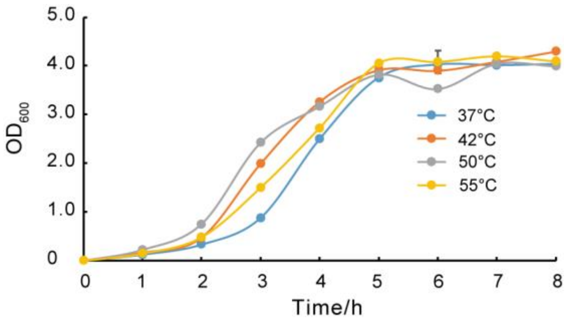


**Supplementary Figure 4. Growth curves of *P. acidilactici* LA412 strain at different temperatures in liquid modified MRS medium.** Data are expressed as the mean ± SD of three technical replicates.

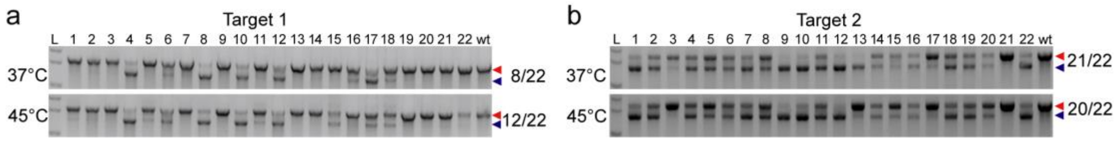


**Supplementary Figure 5. Genome editing in bacterial cells.** Agarose gel analysis of PCR products of the target 1 (**a**) and target 2 (**b**) on *pyrE* gene from randomly selected transformants at 37°C or 45°C. The number on the right of gel bands indicate the editing efficiencies. Red and blue triangles denote wildtype band and deletion band, respectively. L, DNA ladder; 1-22, 22 randomly selected transformants; wt: wildtype cell control.

### Supplementary Table 1: primers and oligonucleotides used in this study

| Primers and nucleotides | Sequence (5′ to 3′) |
|---|---|
| **Primers for SisTnpB1 cloning and expression** | |
| SiRe0632-F-Nde I | CGCCATATGCATCACCATCACCACCATCATCACTGGAGAGCCAAGGAGAAGAATG |
| SiRe0632-R-Sal I | CGCGTCGACGCTATAACACCAGCTTGCTCAGTTA |
| ωRNA-F-Not I | ATTGCGGCCGCTAATACGACTCACTATAGGGTTAAGAAGGACTTGACTTTG |
| ωRNA-R | GTCGACATACTAGGCAATAAAAACCG |
| SOE ωRNA -HDV-F: | TTATTGCCTAGTATGTCGACGGCCGGCATGGTCCCAGCCTCCTCGCTGGCGCCGGCTG |
| SOE ωRNA Xho I+HDV-R: | GGCTCGAGGTCCCATTCGCCATGCCGAAGCATGTTGCCCAGCCGGCGCCAGCGAGGAG |
| TnpB-D187-F | TTGGAATAGCCCTAGGAGTGGATAAACTAG |
| TnpB-D187-R | CACTCCTAGGGCTATTCCAACCACTTTCCC |
| TnpB-E271-F | TATACGTGGCAGATCTTGATGTAAAGGATA |
| TnpB-E271-R | ATCAAGATCTGCCACGTATATCTCATCATA |
| **Oligonucleotides for dsDNA and ssDNA** | |
| DNA-Target-F(5'FAM) | AGCTTTATATTAGAGATAGCATTCATACATACTAGGCAATAAAAACCGTTAAAGCATCC |
| DNA-Target-R(5'FAM) | GGATGCTTTAACGGTTTTTATTGCCTAGTATGTATGAATGCTATCTCTAATATAAAGCT |
| DNA-target-NoTAM-F(5'FAM) | AGCTTTATATTAGAGATAGCATTCATACATACTAGGCAATAAAAACCGAATTTGCATCC |
| DNA-target -NoTAM-R(5'FAM) | GGATGCAAATTCGGTTTTTATTGCCTAGTATGTATGAATGCTATCTCTAATATAAAGCT |
| DNA-NonTarget-F(5'FAM) | CTAGGGCGCGGCTCTCGCTACGGACGCACGCAGCTTACCGCCCCCAATGGCCCTACGAA |
| DNA-NonTarget-R(5'FAM) | TTCGTAGGGCCATTGGGGGCGGTAAGCTGCGTGCGTCCGTAGCGAGAGCCGCGCCCTAG |
| **Primers for construction of target plasmids** | |
| pTarget-F-NdeI | TATGTTTAACGGTTTTTATTGCCTAGTATG |
| pTarget-R-SalI | TCGACATACTAGGCAATAAAAACCGTTAAACA |
| pTarget-F-NoTAM-NdeI | TATGAAATTCGGTTTTTATTGCCTAGTATG |
| pTarget-R-NonTAM-SalI | TCGACATACTAGGCAATAAAAACCGAATTTCA |
| **Primers for seed and TAM sequence variation** | |
| Mut1~5-F | TATGTTTAAGCCAATTTATTGCCTAGTATG |
| Mut1~5-R | TCGACATACTAGGCAATAAATTGGCTTAAACA |
| Mut2-F | TATGTTTAACGGTTAAATATGCCTAGTATG |
| Mut2-R | TCGACATACTAGGCATATTTAACCGTTAAACA |
| Mut3-F | TATGTTTAACGGTTTTTATACGGAAGTATG |
| Mut3-R | TCGACATACTTCCGTATAAAAACCGTTAAACA |

| | |
|---|---|
| Mut4-F | TATGTTTAACGGTTTTTATTGCCTTCATAG |
| Mut4-R | TCGACTATGAAGGCAATAAAAACCGTTAAACA |
| Mut-5~-1-F | TATGAAATTCGGTTTTTATTGCCTAGTATG |
| Mut-5~-1-R | TCGACATACTAGGCAATAAAAACCGAATTTCA |
| TAM-5A-F | TATGATTAACGGTTTTTATTGCCTAGTATG |
| TAM-5A-R | TCGACATACTAGGCAATAAAAACCGTTAATCA |
| TAM-5C-F | TATGCTTAACGGTTTTTATTGCCTAGTATG |
| TAM-5C-R | TCGACATACTAGGCAATAAAAACCGTTAAGCA |
| TAM-5G-F | TATGGTTAACGGTTTTTATTGCCTAGTATG |
| TAM-5G-R | TCGACATACTAGGCAATAAAAACCGTTAACCA |
| TAM-4C-F | TATGTCTAACGGTTTTTATTGCCTAGTATG |
| TAM-4C-R | TCGACATACTAGGCAATAAAAACCGTTAGACA |
| TAM-4G-F | TATGTGTAACGGTTTTTATTGCCTAGTATG |
| TAM-4G-R | TCGACATACTAGGCAATAAAAACCGTTACACA |
| TAM-4A-F | TATGTATAACGGTTTTTATTGCCTAGTATG |
| TAM-4A-R | TCGACATACTAGGCAATAAAAACCGTTATACA |
| TAM-3G-F | TATGTTGAACGGTTTTTATTGCCTAGTATG |
| TAM-3G-R | TCGACATACTAGGCAATAAAAACCGTTCAACA |
| TAM-3C-F | TATGTTCAACGGTTTTTATTGCCTAGTATG |
| TAM-3C-R | TCGACATACTAGGCAATAAAAACCGTTGAACA |
| TAM-3A-F | TATGTTAAACGGTTTTTATTGCCTAGTATG |
| TAM-3A-R | TCGACATACTAGGCAATAAAAACCGTTTAACA |
| TAM-2C-F | TATGTTTCACGGTTTTTATTGCCTAGTATG |
| TAM-2C-R | TCGACATACTAGGCAATAAAAACCGTGAAACA |
| TAM-2G-F | TATGTTTGACGGTTTTTATTGCCTAGTATG |
| TAM-2G-R | TCGACATACTAGGCAATAAAAACCGTCAAACA |
| TAM-2T-F | TATGTTTTACGGTTTTTATTGCCTAGTATG |
| TAM-2T-R | TCGACATACTAGGCAATAAAAACCGTAAAACA |
| TAM-1C-F | TATGTTTACCGGTTTTTATTGCCTAGTATG |
| TAM-1C-R： | TCGACATACTAGGCAATAAAAACCGGTAAACA |
| TAM-1G-F： | TATGTTTAGCGGTTTTTATTGCCTAGTATG |
| TAM-1G-R： | TCGACATACTAGGCAATAAAAACCGCTAAACA |
| TAM-1T-F | TATGTTTATCGGTTTTTATTGCCTAGTATG |
| TAM-1T-R | TCGACATACTAGGCAATAAAAACCGATAAACA |
| Mut1-F | TATGTTTAAGGGTTTTTATTGCCTAGTATG |
| Mut1-R | TCGACATACTAGGCAATAAAAACCCTTAAACA |
| Mut2-F | TATGTTTAACCGTTTTTATTGCCTAGTATG |
| Mut2-R | TCGACATACTAGGCAATAAAAACGGTTAAACA |
| Mut3-F | TATGTTTAACGCTTTTTATTGCCTAGTATG |
| Mut3-R | TCGACATACTAGGCAATAAAAAGCGTTAAACA |
| Mut4-F | TATGTTTAACGGATTTTATTGCCTAGTATG |
| Mut4-R | TCGACATACTAGGCAATAAAATCCGTTAAACA |
| Mut5-F | TATGTTTAACGGTATTTATTGCCTAGTATG |

| | |
|---|---|
| Mut5-R | TCGACATACTAGGCAATAAATACCGTTAAACA |
| Mut6-F | TATGTTTAACGGTTATTATTGCCTAGTATG |
| Mut6-R: | TCGACATACTAGGCAATAATAACCGTTAAACA |
| Mut7-F | TATGTTTAACGGTTTATATTGCCTAGTATG |
| Mut7-R | TCGACATACTAGGCAATATAAACCGTTAAACA |
| Mut8-F | TATGTTTAACGGTTTTAATTGCCTAGTATG |
| Mut8-R | TCGACATACTAGGCAATTAAAACCGTTAAACA |
| Mut9-F | TATGTTTAACGGTTTTTTTTGCCTAGTATG |
| Mut9-R | TCGACATACTAGGCAAAAAAAACCGTTAAACA |
| Mut10-F | TATGTTTAACGGTTTTTAATGCCTAGTATG |
| Mut10-R | TCGACATACTAGGCATTAAAAACCGTTAAACA |
| **Primers for DNA targeting and genome editing in bacterial cells** | |
| pMG36e-0632-F | ATCCTCTTCATCCTCTTCGTCTTGGCCCCTTCAGCTTGAGCTCGT |
| pMG36e-0632-R | CGATCGACCCATATTTAAAAAGCTAGCTATAACACCAGCTTGCTC |
| PlacZ-SOE36e-F | ATCCTCTTCATCCTCTTCGTCTTGGCCCCTTCAGCTTGAGCTCGT |
| PlacZ-SOE0632-R | TGGTGGTGATGGTGATGCATAATATCCATTCCCTTCATTT |
| pMG36e-inverse-TEDA-0632-F | TAGCTTTTTAAATATGGGTCGATCG |
| pMG36e-inverse-TEDA-0632-R | CCAAGACGAAGAGGATGAAGAGGAT |
| pMG36e-inverse-TEDA-ωRNA-F | GCGAATGGGACTGCAGTGCATGAGAGCAAAAAAGAGGAGCCG |
| pMG36e-inverse-TEDA-ωRNA-R | GTCCTTCTTAATCTAGAGCATCTAGAGGATCGATCCCCGGGC |
| ωRNA-F | TGCTCTAGATTAAGAAGGAC |
| SOEguide1-ωRNA-R | CCGATCACCGTTGCCGTCGGGTGAACAACCCCGCCCTTAC |
| SOEguide1+HDV-F | CCGACGGCAACGGTGATCGGGGCCGGCATGGTCCCAGCCTCCTCGCTGGCGCCGGCTGG |
| HDV-R | TGCACTGCAGTCCCATTCGCCATGCCGAAGCATGTTGCCCAGCCGGCGCCAGCGAGGAG |
| SOEguide2-ωRNA-R | TTGCGTAGCTAAATGATCAAGTGAACAACCCCGCCCTTAC |
| SOEguide2+HDV-F | TTGATCATTTAGCTACGCAAGGCCGGCATGGTCCCAGCCTCCTCGCTGGCGCCGGCTGG |
| pyrEdonor-Left-F | GGCTGGGGGGCATGGACGTC |
| pyrEdonor-Right-R | TTTCAGACTTTGCAAGCTTACCCGCAAAGTAGACTTGTG |
| guide@native plasmid-F | CGCCCTAAGAAGCAATGAGAGGCCGGCATGGTCCCAGCCT |
| guide@native plasmid-R | TCTCATTGCTTCTTAGGGCGGTGAACAACCCCGCCCTTACGGACG |
| F1 | TTATTCGGGTTGTTTGGCCG |
| R1 | TTGGACTTTTCCACAGCCCA |
| F2 | TTAATGACGTAAATTAAGCAAGAAGATCTCG |
| R2 | ATGAGCACCACTATTTTATCATTCCA |

| | |
|---|---|
| F3 | TTGGCAGTTAAGAGTATTGGATCAGG |
| R3 | TCACAAACATTGAACATTTTTAGCCG |
| pyrE-verification-F | TGAGCAGTTACAAAACGCGCT |
| pyrE-verification-R | GGCCGATAAGTACCGTAGTCAGC |
